# An optimized ChIP-Seq framework for profiling of histone modifications in *Chromochloris zofingiensis*

**DOI:** 10.1101/2021.11.11.468265

**Authors:** Daniela Strenkert, Matthew Mingay, Stefan Schmollinger, Cindy Chen, Ronan C. O’Malley, Sabeeha S. Merchant

## Abstract

The eukaryotic green alga *Chromochloris zofingiensis* is a reference organism for studying carbon partitioning and a promising candidate for the production of biofuel precursors. Recent transcriptome profiling transformed our understanding of its biology and generally algal biology, but epigenetic regulation remains understudied and represents a fundamental gap in our understanding of algal gene expression. Chromatin Immunoprecipitation followed by deep sequencing (ChIP-Seq) is a powerful tool for the discovery of such mechanisms, by identifying genome-wide histone modification patterns and transcription factor-binding sites alike. Here, we established a ChIP-Seq framework for *Chr. zofingiensis* yielding over 20 million high quality reads per sample. The most critical steps in a ChIP experiment were optimized, including DNA shearing to obtain an average DNA fragment size of 250 bp and assessment of the recommended formaldehyde concentration for optimal DNA-protein crosslinking. We used this ChIP-Seq framework to generate a genome-wide map of the H3K4me3 distribution pattern and to integrate these data with matching RNA-Seq data. In line with observations from other organisms, H3K4me3 marks predominantly transcription start sites of genes. Our H3K4me3 ChIP-Seq data will pave the way for improved genome structural annotation in the emerging reference alga *Chr. zofingiensis*.

## INTRODUCTION

Chromatin is the higher order structure in which eukaryotic DNA is organized, facilitating condensed packaging of genomic DNA into the nucleus. Within chromatin, nucleosomes are considered to be the major structural subunit, in which approximately 146 bp of DNA are wrapped around dimers of core histones H2A, H2B, H3 and H4 (Luger, 2003). In addition to its packaging function, histones are hotspots for a variety of different posttranslational modifications (PTMs) that directly affect gene expression and thus contribute to the role of chromatin towards gene regulation (Li et al., 2007; Workman and Kingston, 1998). Modified N-terminal tails of histones emerge from the nucleosome core and are thus accessible to chromatin effectors, enzymes that convert them into regulatory function. Each organisms’ specific histone code consists of both various different modifications (acetylation, mono/di/tri-methylation, phosphorylation, sumoylation or ubiquitination) and also the multitude of possible combinations between them. Accordingly, the histone code provides an additional regulatory layer extending the information content of the genetic code (Luger, 2003). Previous studies have demonstrated that histone lysine methylation plays an important role during active transcription, for reviews see (Martin and Zhang, 2005; Ruthenburg et al., 2007).

In *Sacharomyces cerevisae*, tri-methylation of Lysine 4 at histone H3 (H3K4me3) is a signature motif marking transcription start sites (TSS) of genes that are either actively transcribed or that are poised for transcription (Bernstein et al., 2002; Ng et al., 2003; Santos-Rosa et al., 2002; Schneider et al., 2004). In addition, H3K4me3 was implied in mediating epigenetic memory (Ng et al., 2003). Notably, enrichment of H3K4me3 at TSSs seems to be a highly conserved feature in many taxa. In human cell lines, mouse and fly, TSSs of actively transcribed genes are also marked with H3K4me3 (Barski et al., 2007; Bernstein et al., 2005; Guenther et al., 2007; Schübeler et al., 2004). The same is true in land plants: in Arabidopsis, rice and maize, tri-methylation at lysine 4 of histone H3 occurred predominantly at gene promoters (He et al., 2010; Oh et al., 2008; Wang et al., 2009; Zhang et al., 2009; Zong et al., 2013).

Chromatin immunoprecipitation followed by deep sequencing (ChIP-Seq) is the method of choice for studying histone modification dynamics. ChIP is based on the recovery of DNA by immuno-precipitation using a specific antibody against the DNA-bound protein of interest, such as a modified histone. ChIP’ed DNA may be quantified using (q)PCR, microarrays or deep sequencing. Detailed and optimized ChIP protocols have been published for Tetrahymena (Dedon et al., 1991), Drosophila (Orlando et al., 1997), yeast (Hecht and Grunstein, 1999), mammalian cell lines (Das et al., 2004), Chlamydomonas (Strenkert et al., 2011) and land plants like Arabidopsis (Bowler et al., 2004), maize (Haring et al., 2007) and tomato (Ricardi et al., 2010). Here, we describe an optimized ChIP protocol for the green alga Chromochloris. As illustrated by ChIP-Seq with an antibody targeting H3K4me3, we show that our optimized protocol is suited for ChIP-Seq analyses of histone modifications with proper resolution and high coverage yielding excellent signal to noise during subsequent data analyses.

## RESULTS

### An optimized ChIP-Seq frame work for Chromochloris – an overview of the most critical parameters

Chromatin immunoprecipitation (ChIP) involves cross-linking of the chromatin-bound proteins by formaldehyde, followed by sonication to obtain small DNA fragments. Immunoprecipitation of cross-linked, fragmented material is then carried out using specific antibodies against the DNA-binding protein of interest. As pointed out by us and others, some parameters are crucial for a successful ChIP experiment and need to be individually adjusted for each respective organism (Das et al., 2004). Accordingly, the optimization of the most critical steps for a ChIP protocol in Chromochloris is outlined in more detail below.

In order to be able to identify the location of the histone-bound DNA sequence of interest with proper resolution, it is critical to break the DNA to an average fragment size of around 250 bp (approximate average nucleosome spacing). The hydrodynamic shearing method using sonication is commonly used for this purpose (Orlando et al., 1997; Spencer et al., 2003). A total of three sonication times were tested on Chromochloris samples: 2, 6 and 10×10 seconds. While intact genomic DNA was still visible after a sonication time of 2×10 seconds, sonication for 10×10 seconds yielded the desired, average DNA fragment size of ∼250 (Figure 1A).

**Figure 1.**
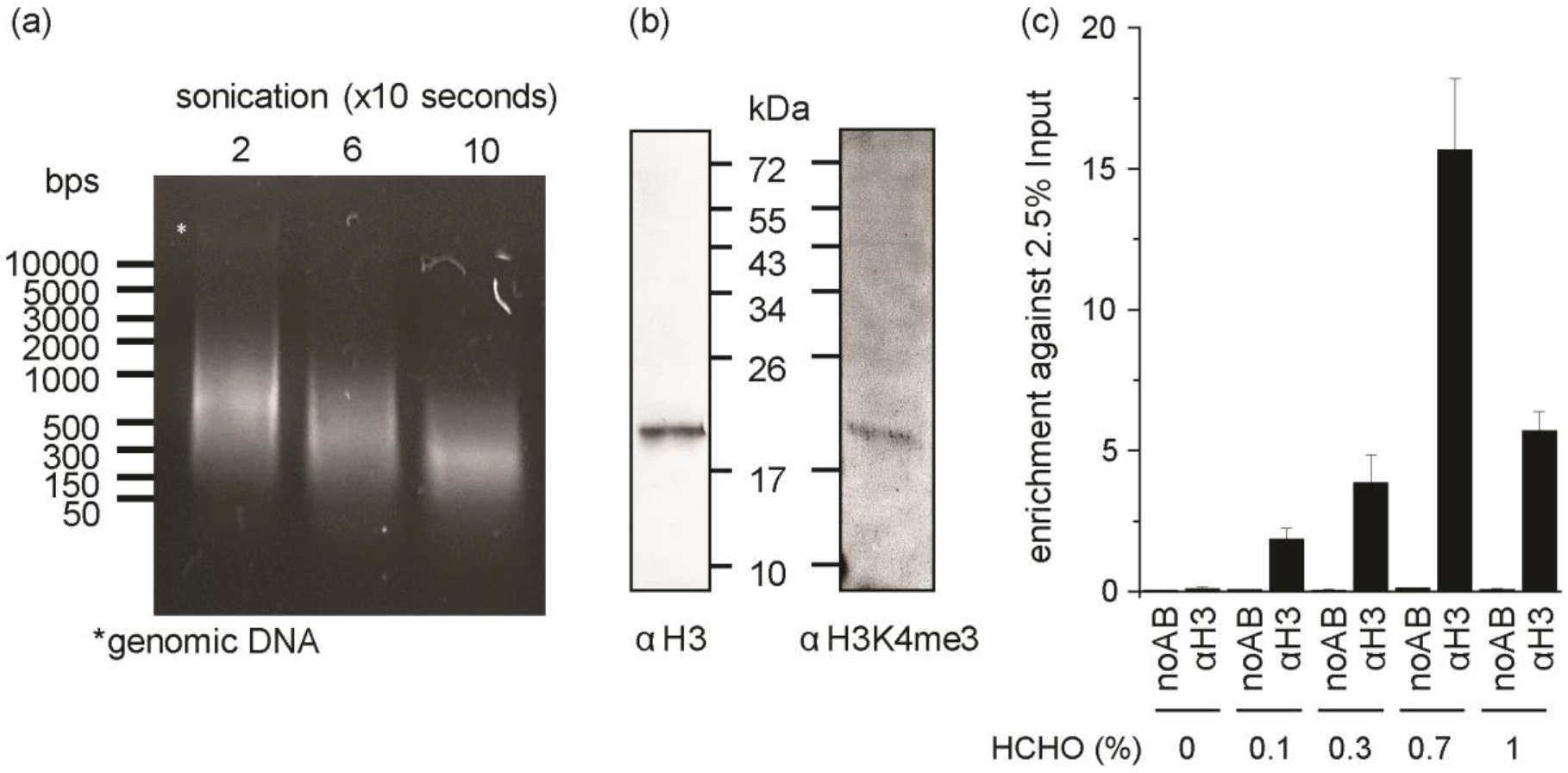
Antibody test, optimization of sonication conditions to yield ∼250-bp chromatin fragments and of formaldehyde concentrations for efficient crosslinking. (a) 2 × 107 cells of *Chromochloris (Chr*.*) zofingiensis* were sonicated 2, 6 and 20 times for 10 seconds. DNA was extracted with phenol/chloroform/isoamyl alcohol (25/24/1 v/v), separated on a 1.5% agarose gel and stained with SYBR gold. (b) Whole-cell proteins corresponding to 2 × 105 cells from Chr. *zofingiensis* were separated on an SDS-polyacrylamide (15% monomer) gel and analyzed by immuno-detection using antibodies against histone H3 or histone H3K4me3 as indicated. (c) 2 × 107 cells were incubated for 10 min with formaldehyde (HCHO) at the concentrations indicated. The reaction was quenched by the addition of 125 mM glycine. Antibodies against histone H3 were used for ChIP (with no antibody as a background control). Chromatin-immunoprecipitated DNA was extracted and amplified using primers targeting the *RBCS* promoter region.

A second important factor for a successful ChIP experiment is the choice of the antibody against the protein of interest. Commercially available antibodies are usually raised against human (modified) histones and are most likely cross-reactive with other species, as histones are highly conserved proteins. Unsurprisingly, Chromochloris histone H3 shares high sequence similarity with human histone H3, making it likely that commercial antibodies will be functional in our applications (Supplemental Figure 1). We aimed to use two different antibodies in this study, one cross-reactive against unmodified histone H3 (for optimization purposes), the other one against tri-methylated H3K4. Cross-reactivity and specificity of both antibodies was tested by immuno-blotting using Chromochloris cell lysates for both of which we saw a single, clear band at the expected molecular weight (Figure 1B).

Crosslinking of the starting material prior to ChIP is used to ensure that the chromatin structure is preserved during sample collection and during the remainder of the Chromatin-immunoprecipitation procedure (Solomon et al., 1988; Solomon and Varshavsky, 1985). Formaldehyde is the most commonly used cross-linking agent in ChIP experiments as it penetrates most cell walls and because cross-links are reversible by incubation at higher temperatures; commonly 65 °C is used for the purpose of de-crosslinking (Orlando et al., 1997). In addition, formaldehyde forms bonds that span a distance of approximately 2 Å (Dedon et al., 1991), resulting in DNA-protein, RNA-protein, as well as protein-protein cross-linking. While one needs sufficient amounts of formaldehyde for efficient DNA-protein crosslinking, it is also crucial not to over-crosslink the sample to avoid the loss of antibody recognition sites and thus loss of ChIP’ed DNA. Many published ChIP protocols use a standard amount of 1% formaldehyde, but we have seen in the past that different concentrations can give better signal to noise ratios in ChIP experiments of algae (Strenkert et al., 2011). For the determination of the optimal crosslinker concentrations, ChIP was performed using an antibody targeting unmodified histone H3 (as opposed to a certain histone mark) as we expected reasonable nucleosome occupancy at the whole-genome level, including at a promoter region of the gene encoding the small subunit of Rubisco, *RBCS*. We tested formaldehyde at concentrations of 0, 0.1, 0.3, 0.7 and 1 percent. Samples without an antibody (noAB) were used as negative controls, which allowed us to estimate DNA enrichment over background contamination if histone-DNA crosslinking was successful. While all formaldehyde concentrations tested led to significant enrichment of ChIP’ed DNA over background (Figure 2C), the use of 0.7% formaldehyde yielded the largest amounts of ChIP’ed RBCS promoter DNA and the highest signal to noise (Figure 2C). Lower formaldehyde concentrations gave insufficient crosslinking, and higher concentrations resulted in over-crosslinking, both of which reduced ChIP efficiency. Therefore, we recommend the use of 0.7% formaldehyde.

**Figure 2.**
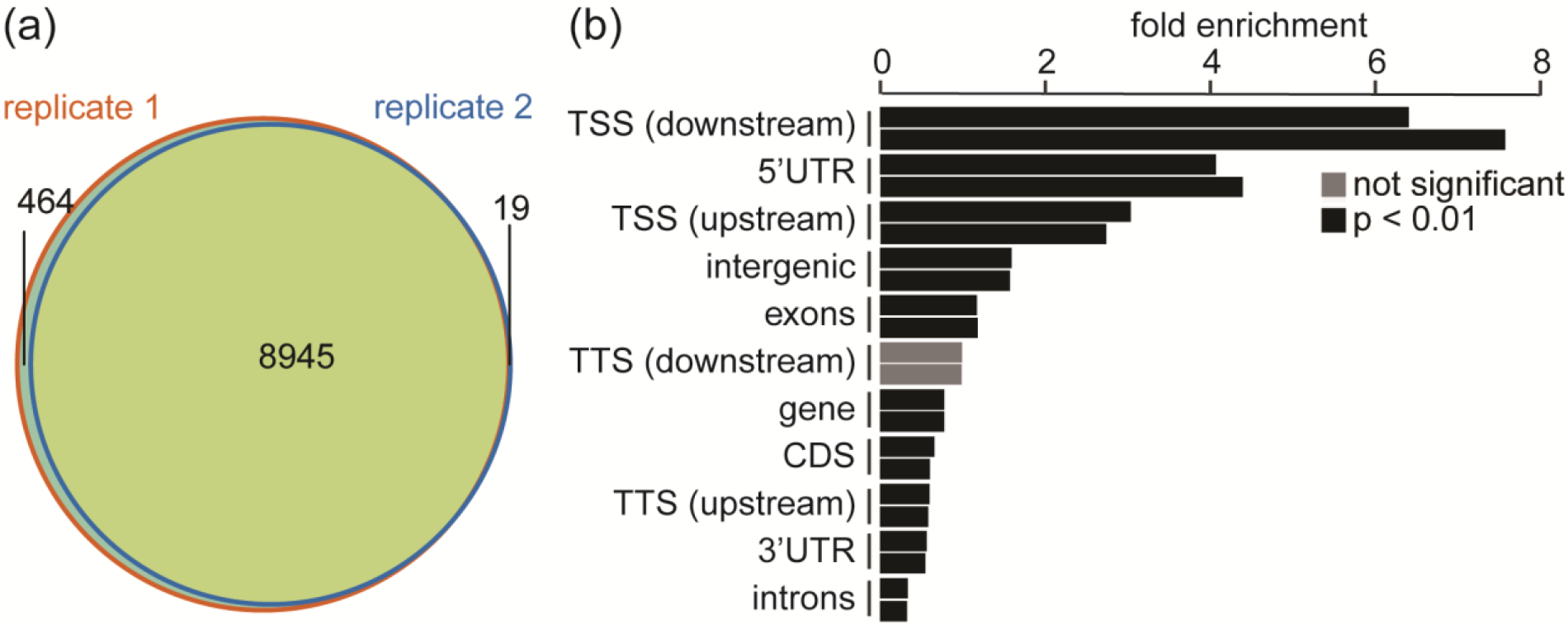
H3K4me3 peak calling and genomic distribution in Chromochloris. (a) Total number of narrow H3K4me3 peaks in each replicate as well as the overlap between both replications. (b) Relative fold-enrichment of H3K4me3 at different genomic regions as indicated. Each individual sample represents one independent replicate. The fold-enrichment represents the number of peaks that overlap with a region divided by the number of peaks that would overlap with the respective region if they were randomly distributed across the genome.

### A survey of the genome wide H3K4me3 distribution pattern in Chromochloris

To further test suitability of our optimized ChIP framework for deep sequencing applications, we determined H3K4me3 enrichment at a genome-wide level. H3K4me3 is a so-called narrow-peak histone mark that seems to be specifically enriched at TSSs of protein coding genes in a variety of different organisms including yeast, mammals and land plants (Barski et al., 2007; Bernstein et al., 2005; Pokholok et al., 2005; Santos-Rosa et al., 2002; Zhang et al., 2009). Accordingly, assessment of data quality will be straightforward. In our ChIP experiments, one possible negative control, ChIPs performed without an antibody (noAB controls) did not yield sufficient amounts of DNA for library preparation and subsequent deep sequencing. It is certainly possible to identify DNA binding sites based on relative enrichment of different chromosomal regions even without a negative control, however, some chromosomal regions can be significantly enriched even when sequencing genomic DNA alone (Rozowsky et al., 2009). Accordingly, non-ChIP’ed, genomic DNA is often analyzed in parallel as negative control for ChIP-Seq experiments to account for this issue. To this end, we included non-ChIP’d genomic DNA as our control. We interrogated H3K4 tri-methylation from duplicate samples in Chromochloris cultures. mRNA from the same cultures was obtained in parallel and used for RNA-Seq analysis, with the intention to later analyze it alongside the ChIP-Seq data. By aligning H3K4me3 ChIP-Seq reads to the reference genome, we noted that the majority of reads (> 98%) could be mapped to the nuclear genome, which suggested that the sequencing data were of a high quality (Table 1). Model-based analysis using ChIP-Seq (MACS) software (Zhang et al., 2008) was used for peak calling. We identified a total of 8945 H3K4me3 consensus peaks in both replicates, with the duplicate samples showing excellent overlap between each other (Figure 2A).

**Table 1.**
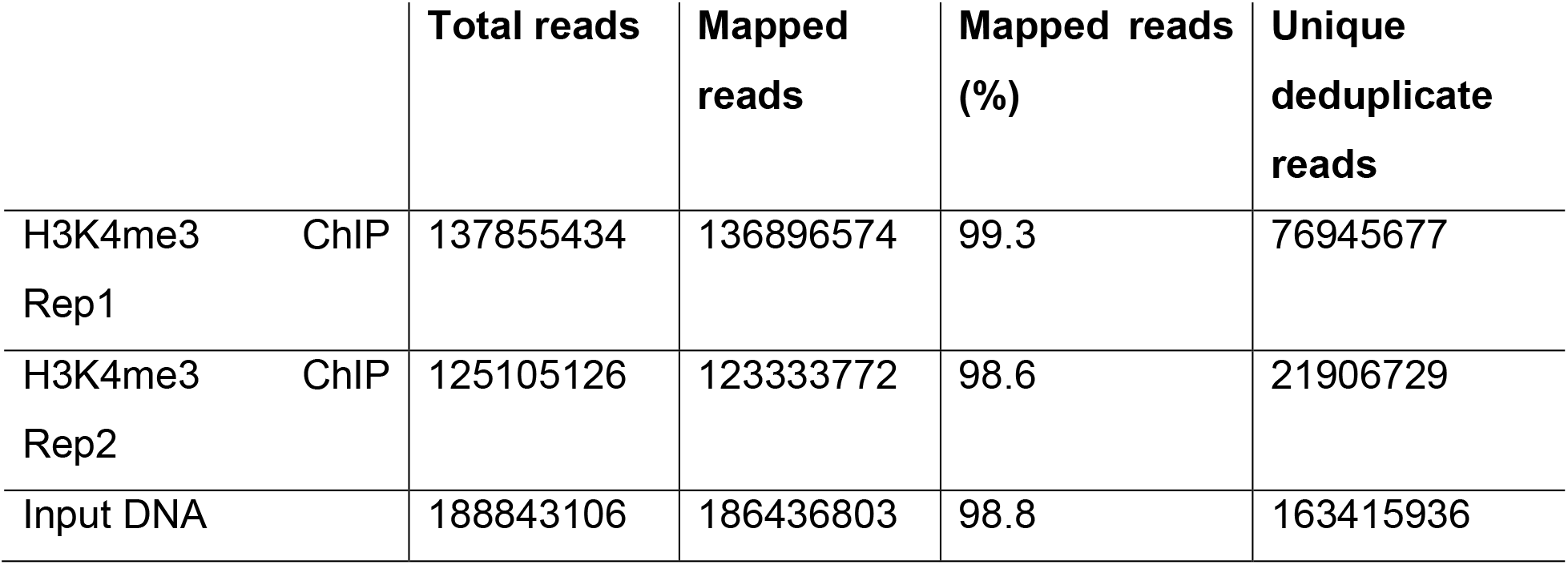
H3K4me3 ChIP-seq mapping statistics.

The distribution of H3K4me3 peaks along the Chromochloris genome was further characterized using eleven groups that included distinct genic regions and observed predominant H3K4me3 enrichment at TSSs of genes (Figure 2B). Genome-wide profiling of H3K4me3 revealed an average H3K4me3 peak width of +/-500 bp around the TSSs in the majority of genes (Figure 3).

**Figure 3.**
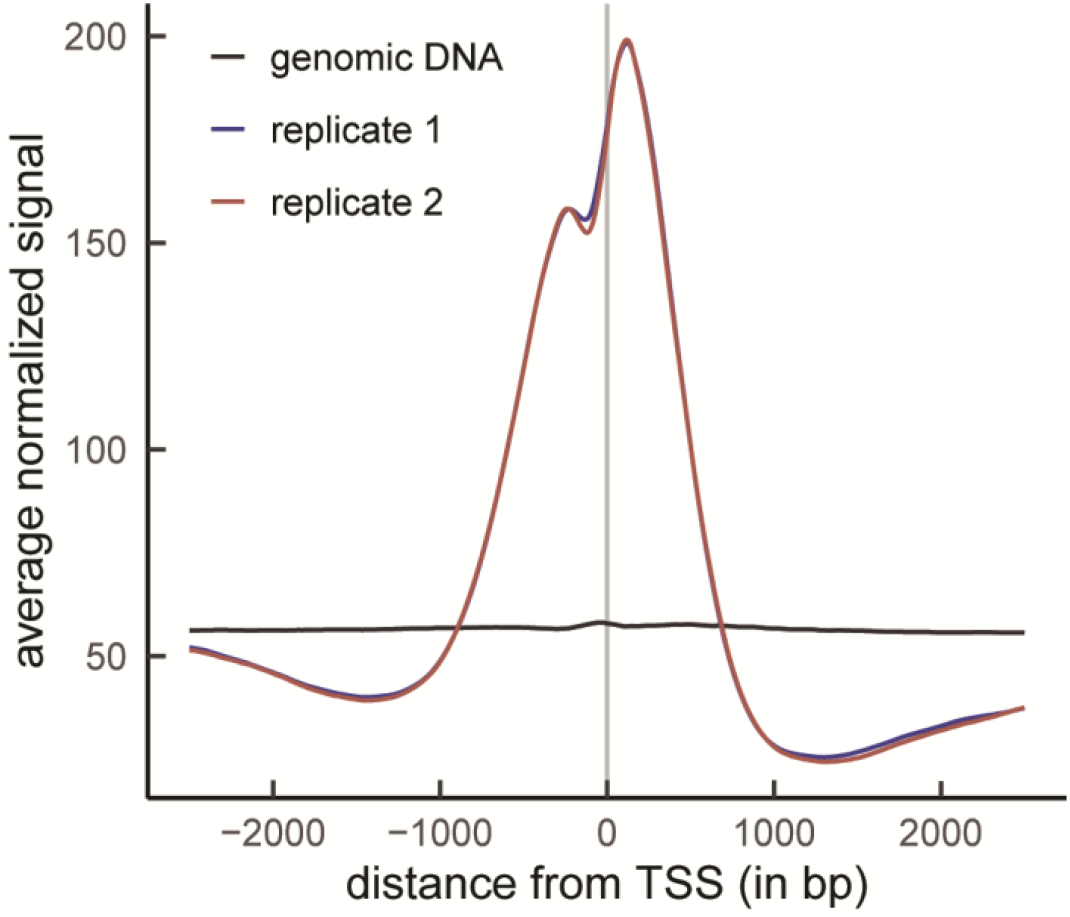
Gene wide distribution pattern of H3K4me3 in Chromochloris. Shown is the average normalized H3K4me3 enrichment for both replicates (blue and red) and input genomic DNA (black) at the TSSs (grey line) and +/-2500 bps from TSSs. Values were normalized by scaling based on the number of uniquely mapped, non-duplicate reads.

### H3K4me3 and gene expression

To get an integrated view on H3K4me3 enrichment and its relationship to gene expression, we compared transcript abundances of individual genes with H3K4me3 enrichment at their respective TSSs. An increase in transcript abundance seems to correlate with the chances of a gene harboring a H3K4me3 at its TSS (Figure 4). When we further interrogated transcript abundances of genes that harbor a H3K4me3 peak at their TSS and compared it to those that do not, again, we saw a clear, positive correlation between active transcription and the likelihood of a gene being marked with H3K4me3 (Figure 5).

**Figure 4.**
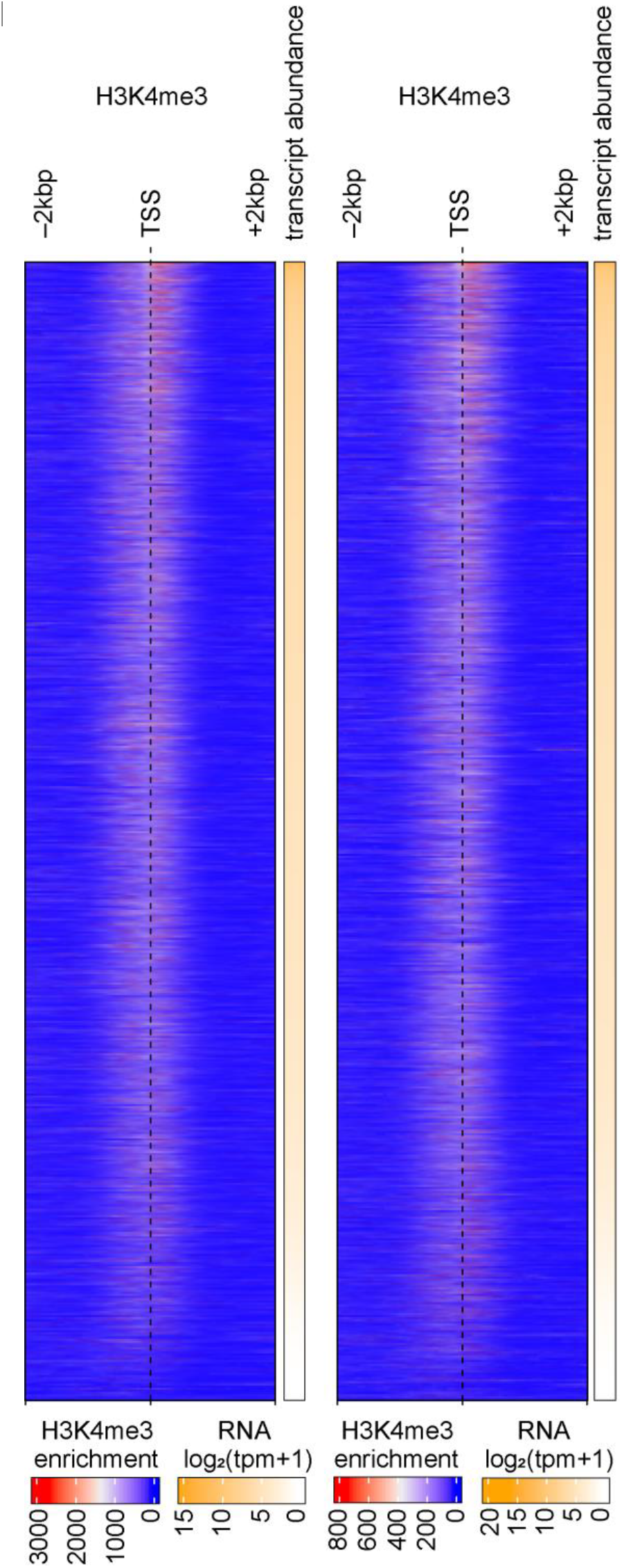
H3K4me3 enrichment ranked by gene expression. Shown are H3K4me3 values visualized as heat map. H3K4me3 values are sorted based on the respective expression values (as Transcripts Per Million, TPMs) for both replicates at the TSSs (dashed line) and +/-2000 bp from TSSs.

**Figure 5.**
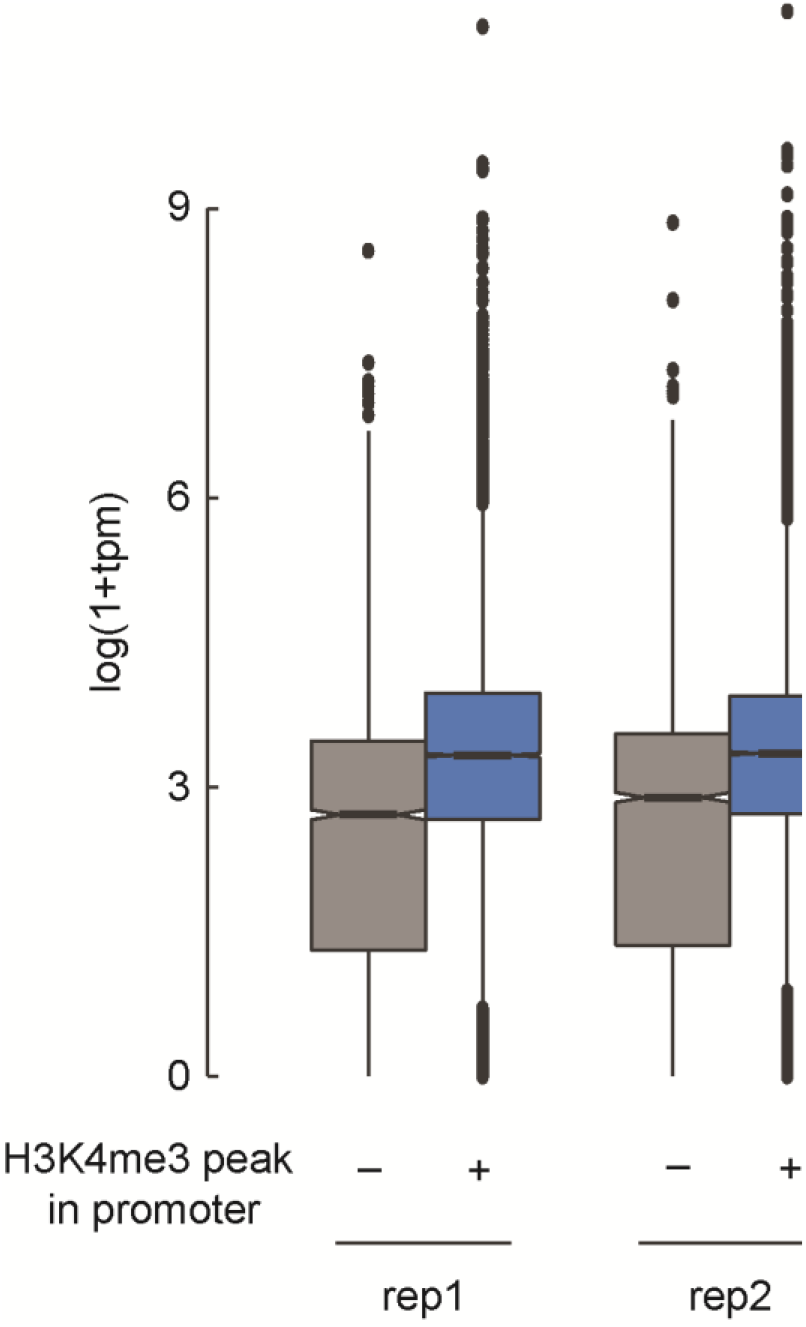
Integrated analyses of H3K4me3 enrichment and gene expression. Transcript abundance values (in TPMs -Transcripts Per Kilobase Million) from genes that harbor a H3K4me3 peak at their TSSs (blue) vs all others (grey) are visualized using boxplots.

## DISCUSSION

Epigenetic regulation is a prerequisite for mediating transcriptional responses in general and for the fine tuning of genetically programmed changes in particular. Chromatin immunoprecipitation coupled with deep sequencing (ChIP-Seq) is a powerful tool to assess changes in histone mark patterns and to identify transcription factor binding sites. Most importantly, some critical parameters in the procedure need to be optimized in each organism individually to ensure proper data quality and reproducibility between samples. Given the conserved, genome wide distribution pattern of H3K4me3 in many diverse taxa, ChIP-Seq using H3K4me3 was facilitated as a proof of principle application to illustrate desired data quality of our optimized ChIP-Seq frame work. H3K4me3 was shown to be a narrow-peak histone mark that is enriched at TSSs of genes in yeast, fly, human, algae and plants (Barski et al., 2007; Bernstein et al., 2005; Pokholok et al., 2005; Santos-Rosa et al., 2002; Zhang et al., 2009). Unsurprisingly, and in line with a conserved role for H3K4me3 in gene activation, Chromochloris genes are also marked with H3K4me3 at their TSSs. Notably, the likelihood of a gene being marked by H3K4me3 was higher, if the underlying gene was transcribed.

The optimized ChIP-Seq frame-work established herein will pave the way for studying epigenetic changes and transcription factor binding sites in the emerging reference alga Chromochloris. Moreover, the coverage achieved by our ChIP-Seq frame work will be sufficient for quantitative recovery of narrow and broad peak histone marks alike. This work represents a critical first step in establishing chromatin state dynamics in Chromochloris.

## MATERIALS AND METHODS

### Cultures and growth conditions

*Chromochloris zofingiensis* SAG 211-14 culture was grown in TAP medium with KNO_3_ replacing NH_4_Cl as nitrogen source and with modified trace elements from (Kropat et al., 2011) instead of Hutner’s trace elements. Cells were grown at 90 µmol photons m^-2^ s^-1^, 140 rpm to a cell density of 2 × 10^6^ cells/mL for all experiments.

### Cross-reactivity and specificity of antibodies used for ChIP

All antibodies used in this study were evaluated on total cell lysate of Chromochloris SAG 211-14 cultures. 2 ×10^7^ cells were collected by centrifugation at 4°C, 1650 × *g* and 2 min. The cell pellet was resuspended in 500 µL MilliQ water and 2x protein lysis buffer was added (125 mM Tris-Cl pH 6.8, 20% glycerol, 4% SDS, 10% β-mercaptoethanol and 0.005% bromophenol blue). Samples were sonicated for 5 seconds to break the cell wall (Sonic Dismembrator System with a 1/2 in. Probe, Model 505; 117V, 50/60Hz (Fisher Scientific Company, No.: FB505110), settings: 1 sec ON / 1 sec OFF, Amplitude 50%). 20 µL cell lysate corresponding to 4 × 10^5^ cells was loaded on each lane. Proteins were separated by 15% SDS-PAGE and transferred to nitrocellulose membranes (Amersham Protran 0.1 NC). The membrane was blocked for 30 min with 3% dried non-fat milk in phosphate buffered saline (PBS) solution containing 0.1% (w/v) Tween 20 and then incubated in primary antiserum. The PBS solution was used as the diluent for both primary and secondary antibodies. The membranes were washed in PBS containing 0.1% (w/v) Tween 20. Antibodies directed against histone H3 (1:1000) or tri-methylated lysine 4 of histone H3 (1:1000) were used. The secondary antibody, used at 1:4000, was goat anti-rabbit conjugated to alkaline phosphatase, and processed according to the manufacturer’s instructions.

### Establishment of DNA shearing

A culture volume corresponding to 2 ×10^8^ was collected by centrifugation at 4°C, 1650 × *g* for 2 minutes. Cell pellet was resuspended in 1 mL ChIP lysis buffer (1% SDS, 10 mM EDTA, 50 mM Tris-Cl, pH 8.0, and 0.25 x protease inhibitor cocktail [Roche]) and transferred to Beckman Coulter 4mL Polycarbonate Thick Wall Centrifuge Tubes (13 × 64 mm). For shearing of genomic DNA, we used a Sonic Dismembrator System with a 1/2 in. Probe, Model 505; 117V, 50/60Hz (Fisher Scientific Company, No.: FB505110) settings: 1 sec ON / 1 sec OFF, Amplitude 50%). To test conditions needed for efficient fragmentation of Chromochloris genomic DNA, we used 2, 6 and 10 seconds of sonication. After sonication, 100 µL of each cell lysate (corresponding to 2 × 10^7^ cells) was transferred to a new 1.5 mL Eppendorf tube. 300 µL of ChIP lysis buffer was added. DNA was extracted using phenol/chloroform extraction as follows: 500 µL of phenol/chloroform/Isoamyl alcohol (25:24:1) was added to the cell lysates, samples were mixed and centrifuged for 10 minutes at 4°C and 16200 × *g*. The aqueous phase was transferred to 500 µL chloroform/isosamyl alcohol (24:1). Samples were mixed and centrifuged for 10 minutes at 4 °C and 16200 x *g*, this step was repeated once. After centrifugation, the aqueous phase containing the DNA was added to a new Eppendorf tube containing 50 µL 3M Na-acetate (pH 5.3) and 1 mL of 100% ethanol was added. Samples were mixed and kept at -20°C for at least 2 hours. DNA was pelleted by a 15 min centrifugation at 4°C at 16200 x *g*. DNA pellets were washed once with 70% ethanol, and the DNA pellet was air dried for at least 15 minutes at room temperature. DNA was resuspended in 50 µL TE supplemented with 10 µg/µL DNase-free RNAse A. 10 µL of each sample was supplemented with 6 × DNA loading dye and loaded on a 1.2% Agarose gel supplemented with SYBR gold.

### Optimizing Formaldehyde crosslinking

A culture volume corresponding to 2 × 10^8^ was collected by centrifugation at 4°C, 1650 x *g* for 2 minutes. Cell pellets were washed with 50 mL KH buffer (20 mM K-HEPES, pH 7.4, 40 mM KCl). This is necessary because residual Tris from the growth medium would quench formaldehyde and lead to irreproducible results. After washing, cells were collected by centrifugation for 2 minutes at 4°C and 1650 x *g*. Cell pellets were resuspended in 10 mL crosslinking buffer (20 mM K-HEPES, 40 mM KCl supplemented with no or 0,1, 0.3, 0.7 and 1% formaldehyde). This solution was always made fresh because formaldehyde is unstable. Optimal formaldehyde concentration for crosslinking DNA to chromatin in Chromochloris was determined based on efficient Chromatin immunoprecipitation (ChIP) using an antibody cross-reactive against an unmodified N-terminus of Histone H3. We used the promoter region of *RBCS* as target genomic region during optimization. qPCR was performed on a Bio-Rad CFX96 Touch Real-Time PCR Detection System using iTAQ Mastermix according to the manufacturer’s instructions. Primers used were as follows: RBCSpromfor: CAATGCAAGCAGTTCGCATG and RBCSpromrev: ACGGAGGACTTGGCAATGAC.

### Optimized Chromatin-immunoprecipitation (ChIP) protocol

A culture volume corresponding to 2 × 10^8^ cells was collected by centrifugation at 4°C, 1650 x *g* for 2 minutes. Cell pellets were washed with 50 mL KH buffer. After washing, cells were collected by centrifugation for 2 minutes at 4°C and 1650 x *g*. Cell pellets were resuspended in 10 mL crosslinking buffer (20 mM HEPES-KOH, 40 mM KCL supplemented with 0.7 % formaldehyde). The cell pellet was resuspended in 1 mL ChIP lysis buffer (1% SDS, 10 mM EDTA, 50 mM Tris-Cl, pH 8.0, and 0.25 x protease inhibitor cocktail [Roche]) and transferred to Beckman Coulter 4mL Polycarbonate Thick Wall Centrifuge Tubes (13 × 64 mm). For shearing of genomic DNA, we used a Sonic Dismembrator System with a 1/2 in. probe Model 505; 117V, 50/60Hz (Fisher Scientific Company, No.: FB505110) at settings: 1 sec ON / 1 sec OFF, Amplitude 50%). After sonication for 10 seconds, cell lysate was centrifuged for 10 mins at 4°C and 16200 x *g* (this is to clear the lysate from starch and cell debris). We generated aliquots of 100 μl Input chromatin in 1.5 mL tubes and flash froze them in liquid nitrogen (each aliquot corresponds to chromatin from ∼2 × 10^7^ cells). The input chromatin can be stored for several months at -80°C, but avoid multiple freeze/thaw cycles. We used 50 mg of Protein-A-Sepharose beads (Sigma, P3391-1G), resuspended beads in 1 ml ChIP-buffer (1.1% Triton X-100, 1.2 mM EDTA, 167 mM NaCl, and 16.7 mM Tris-HCl, pH 8) and incubated them for 30 min at 4°C. Swollen beads were washed two times with 500 μl ChIP buffer. After the last wash, supernatant was discarded and we added 500 μl ChIP-buffer (gives ∼750 μl suspension with swollen beads). We did not add sonicated lambda DNA, since this would interfere with DNA library generation. For ChIP, 5 μl of anti-H3 or 10 μl of anti-H3K4me3 antibodies was premixed with 10 μl BSA solution (10 mg/mL) and incubated for at least 30 min on ice. For each antibody employed, two 100-μl aliquots of chromatin solution were thawed on ice (one for a control without antibody and one for the antibody to be tested). 900 μl of ChIP buffer was added to each aliquot (from this step on we consequently used filter tips to avoid DNA contaminations). Samples were centrifuged for 20 sec at 16,100 x *g* and 4°C, and supernatant was transferred to microcentrifuge tubes containing the prepared antibody solutions. Samples were mixed on a rotation wheel for 1 h at 4°C. After 1 hour, samples were centrifuged for 20 sec at 16200 x *g* and 4°C, and supernatants were transferred to new microcentrifuge tubes containing 60 μl Sepharose beads. Samples were mixed on a rotation wheel for 2 h at 4°C. After incubation, samples were centrifuged for 20 sec at 16200 x *g* and 4°C and supernatants discarded. Sepharose beads were washed once with washing buffer 1 (0.1% SDS, 1% Triton X-100, and 2 mM EDTA, pH 8) containing 150 mM NaCl, once with washing buffer 1 containing 500 mM NaCl, once with washing buffer 2 (250 mM LiCl, 1% Nonidet P-40, 1% Na-deoxycholate, 1 mM EDTA, and 10 mM Tris-HCl, pH 8), and twice with TE (1 mM EDTA and 10 mM Tris-HCl, pH 8). Cross-links were reverted by an overnight incubation at 65°C after addition of NaCl to a final concentration of 0.5 M, 1% SDS and 0.1 M NaHCO_3_(made fresh) for 15 min at 65°C. This step was repeated once and eluates were pooled. At this point another aliquot of Input chromatin was also thawed, 400 µL of ChIP Lysis buffer was added and incubated overnight at 65°C (Input DNA control). In order to remove proteins in the precipitates, we did add 10 μl 0.5 M EDTA (pH 8.0), 20 μl 1 M Tris-HCl (pH 8.0) and 2.1 μl proteinase K (10 mg/ml) and incubated the samples for 1 h at 55°C. DNA extraction was performed by one extraction with 500 μl phenol/chloroform/isoamyl alcohol and one extraction using 500 μl chloroform/isoamyl alcohol. DNA was precipitated by adding 50 μl 3 M Na-acetate (pH 5.2), 2.5 μl glycogen (2.5 μg/μl) (this will give an otherwise invisible pellet) and 1 ml 100% EtOH and incubating for at least 2 h at -20°C. Samples were centrifuged for 20 min at 16200 x *g* and 4°C. DNA pellet was dried and resuspended in 50 μl 10mM Tris-HCl buffer (pH 8.0).

### ChIP library preparation and sequencing

5 ng of ChIP’d DNA was treated with end-repair, A-tailing, and ligation of Illumina compatible adapters (IDT) using the KAPA-HyperPrep kit (KAPA biosystems). The ligated products were enriched with 8 cycles of PCR (KAPA biosystems) and size selected to 200-500 bp with Total Pure NGS beads (Omega Bio-tek). The prepared libraries were quantified using KAPA Biosystems’ next-generation sequencing library qPCR kit and run on a Roche LightCycler 480 real-time PCR instrument. Sequencing of the flow cell was performed on the Illumina NextSeq500 sequencer using NextSeq500 NextSeq HO kits, v2, following a 2×151 indexed run recipe.

### ChIP Read preprocessing and filtering

2×151 sequence data was generated at the DOE Joint Genome Institute (JGI) using the Illumina NextSeq platform. BBDuk version 38.87 (http:\\bbtools.jgi.doe.go) was used to remove contaminants, trim reads that contained adapter sequence and homopolymers of G’s of size 5 or more at the ends of the reads and right quality trim reads where quality drops below 6. BBDuk was used to remove reads that contained 1 or more ‘N’ bases and had an average quality score across the read less than 10 or had minimum length <= 49 bp. Reads mapped with BBMap (v. 38.87) to masked human, cat, dog, mouse and common microbial contaminant references at 93% identity were removed.

### ChIP Read Alignment and Peak Calling

Filtered reads were aligned to the *Chromochloris zofingiensis* reference genome (Roth et al., 2017) using BWA mem (version 0.7.17). Only uniquely mapped reads were retained. Duplicated reads were removed by Picard (v. 2.22.9) (http://broadinstitute.github.io/picard/; Broad Institute) MarkDuplicates tool. Finally, the remaining reads were used for peak calling by MACS2 (v. 2.1.1) (Zhang et al., 2008) with parameters “--call-summits --nomodel --extsize 147 -c”. Input control libraries were generated and used for peak calling and downstream analysis. To visualize and plot data, bigwig files were created using bedGraphToBigWig (v.4) (Kent et al., 2010) and Deeptools (v. 3.1.3) (Ramirez et al., 2016) was used to generate summary signal plots and heatmaps.

### RNA extraction, sequencing and transcriptome analyses

A culture volume corresponding to 5 × 10^7^ cells was collected by centrifugation at 4°C, 3500 rpm for 2 mins. Cell pellet was resuspended in 0.2 mL RLC buffer (Qiagen), flash frozen in liquid nitrogen and ground to a fine powder using mortar and pestle. Sample was added to 700 µL Trizol and mixed overhead before the addition of 200 μl Chloroform/Isoamyl alcohol. Samples were vigorously shaken, then centrifuged for 10 mins at 4°C and 13200 rpm. Supernatant was added to 700 μl isopropanol. RNA was precipitated at -20°C over night and washed with 70% Ethanol, air dried and resuspended in 40 μl RNAse free water. DNase I digest and clean up was performed using Zymo according to the manufacturer’s instructions.

RNA was converted into cDNA and made into sequence ready libraries with the KAPA RNA-Seq Kit (Kapa Biosystems). RNA-Seq libraries were sequenced with 150 bp single-end reads on a HiSeq 2500. Data was aligned to the ChrZofV5 release of the C. zofingiensis genome with RNA STAR. Determination of counts per gene and transcript abundance in transcripts per million (TPMs) were made with DESeq2.

## ACKNOWLEDGMENTS

This material is based upon work supported by the U.S. Department of Energy, Office of Science, Office of Biological and Environmental Research, under Award Number DE-SC0018301. We also thank Kris K. Niyogi for critical reading of the manuscript and kindly providing the histone H3 antibody. This research was performed under the Facilities Integrating Collaborations for User Science (FICUS) initiative and used resources at the DOE Joint Genome Institute which is a DOE Office of Science User Facility sponsored by the Office of Biological and Environmental Research and operated under Contract Nos. DE-AC02-05CH11231.

## Notes

### Competing Interest Statement

The authors have declared no competing interest.

## REFERENCES

Barski, A., S. Cuddapah, K. Cui, T.Y. Roh, D.E. Schones, Z. Wang, G. Wei, I. Chepelev, and K. Zhao. 2007. High-resolution profiling of histone methylations in the human genome. Cell. 129:823–837.

Bernstein, B.E., E.L. Humphrey, R.L. Erlich, R. Schneider, P. Bouman, J.S. Liu, T. Kouzarides, and S.L. Schreiber. 2002. Methylation of histone H3 Lys 4 in coding regions of active genes. Proc Natl Acad Sci U S A. 99:8695–8700.

Bernstein, B.E., M. Kamal, K. Lindblad-Toh, S. Bekiranov, D.K. Bailey, D.J. Huebert, S. McMahon, E.K. Karlsson, E.J. Kulbokas, 3rd, T.R. Gingeras, S.L. Schreiber, and E.S. Lander. 2005. Genomic maps and comparative analysis of histone modifications in human and mouse. Cell. 120:169–181.

Bowler, C., G. Benvenuto, P. Laflamme, D. Molino, A.V. Probst, M. Tariq, and J. Paszkowski. 2004. Chromatin techniques for plant cells. Plant J. 39:776–789.

Das, P.M., K. Ramachandran, J. vanWert, and R. Singal. 2004. Chromatin immunoprecipitation assay. BioTechniques. 37:961–969.

Dedon, P.C., J.A. Soults, C.D. Allis, and M.A. Gorovsky. 1991. A simplified formaldehyde fixation and immunoprecipitation technique for studying protein-DNA interactions. Analytical biochemistry. 197:83–90.

Guenther, M.G., S.S. Levine, L.A. Boyer, R. Jaenisch, and R.A. Young. 2007. A chromatin landmark and transcription initiation at most promoters in human cells. Cell. 130:77–88.

Haring, M., S. Offermann, T. Danker, I. Horst, C. Peterhansel, and M. Stam. 2007. Chromatin immunoprecipitation: optimization, quantitative analysis and data normalization. Plant Methods. 3:11.

He, G.M., X.P. Zhu, A.A. Elling, L.B. Chen, X.F. Wang, L. Guo, M.Z. Liang, H. He, H.Y. Zhang, F.F. Chen, Y.J. Qi, R.S. Chen, and X.W. Deng. 2010. Global Epigenetic and Transcriptional Trends among Two Rice Subspecies and Their Reciprocal Hybrids. The Plant cell. 22:17–33.

Hecht, A., and M. Grunstein. 1999. Mapping DNA interaction sites of chromosomal proteins using immunoprecipitation and polymerase chain reaction. Methods in enzymology. 304:399–414.

Kent, W.J., A.S. Zweig, G. Barber, A.S. Hinrichs, and D. Karolchik. 2010. BigWig and BigBed: enabling browsing of large distributed datasets. Bioinformatics. 26:2204–2207.

Kropat, J., A. Hong-Hermesdorf, D. Casero, P. Ent, M. Castruita, M. Pellegrini, S.S. Merchant, and D. Malasarn. 2011. A revised mineral nutrient supplement increases biomass and growth rate in Chlamydomonas reinhardtii. Plant J. 66:770–780.

Li, B., M. Carey, and J.L. Workman. 2007. The role of chromatin during transcription. Cell. 128:707–719.

Luger, K. 2003. Structure and dynamic behavior of nucleosomes. Curr Opin Genet Dev. 13:127–135.

Martin, C., and Y. Zhang. 2005. The diverse functions of histone lysine methylation. Nature reviews. Molecular cell biology. 6:838–849.

Ng, H.H., F. Robert, R.A. Young, and K. Struhl. 2003. Targeted recruitment of Set1 histone methylase by elongating Pol II provides a localized mark and memory of recent transcriptional activity. Molecular cell. 11:709–719.

Oh, S., S. Park, and S. van Nocker. 2008. Genic and global functions for Paf1C in chromatin modification and gene expression in Arabidopsis. PLoS Genet. 4:e1000077.

Orlando, V., H. Strutt, and R. Paro. 1997. Analysis of chromatin structure by in vivo formaldehyde cross-linking. Methods. 11:205–214.

Pokholok, D.K., C.T. Harbison, S. Levine, M. Cole, N.M. Hannett, T.I. Lee, G.W. Bell, K. Walker, P.A. Rolfe, E. Herbolsheimer, J. Zeitlinger, F. Lewitter, D.K. Gifford, and R.A. Young. 2005. Genome-wide map of nucleosome acetylation and methylation in yeast. Cell. 122:517–527.

Ramirez, F., D.P. Ryan, B. Gruning, V. Bhardwaj, F. Kilpert, A.S. Richter, S. Heyne, F. Dundar, and T. Manke. 2016. deepTools2: a next generation web server for deep-sequencing data analysis. Nucleic Acids Res. 44:W160–165.

Ricardi, M.M., R.M. González, and N.D. Iusem. 2010. Protocol: fine-tuning of a Chromatin Immunoprecipitation (ChIP) protocol in tomato. Plant Methods. 6:1746–4811.

Roth, M.S., S.J. Cokus, S.D. Gallaher, A. Walter, D. Lopez, E. Erickson, B. Endelman, D. Westcott, C.A. Larabell, S.S. Merchant, M. Pellegrini, and K.K. Niyogi. 2017. Chromosome-level genome assembly and transcriptome of the green alga Chromochloris zofingiensis illuminates astaxanthin production. P Natl Acad Sci USA. 114:E4296–E4305.

Rozowsky, J., G. Euskirchen, R.K. Auerbach, Z.D.D. Zhang, T. Gibson, R. Bjornson, N. Carriero, M. Snyder, and M.B. Gerstein. 2009. PeakSeq enables systematic scoring of ChIP-seq experiments relative to controls. Nature Biotechnology. 27:66–75.

Ruthenburg, A.J., C.D. Allis, and J. Wysocka. 2007. Methylation of lysine 4 on histone H3: intricacy of writing and reading a single epigenetic mark. Molecular cell. 25:15–30.

Santos-Rosa, H., R. Schneider, A.J. Bannister, J. Sherriff, B.E. Bernstein, N.C. Emre, S.L. Schreiber, J. Mellor, and T. Kouzarides. 2002. Active genes are tri-methylated at K4 of histone H3. Nature. 419:407–411.

Schneider, R., A.J. Bannister, F.A. Myers, A.W. Thorne, C. Crane-Robinson, and T. Kouzarides. 2004. Histone H3 lysine 4 methylation patterns in higher eukaryotic genes. Nat Cell Biol. 6:73–77.

Schübeler, D., D.M. MacAlpine, D. Scalzo, C. Wirbelauer, C. Kooperberg, F. van Leeuwen, D.E. Gottschling, L.P. O’Neill, B.M. Turner, J. Delrow, S.P. Bell, and M. Groudine. 2004. The histone modification pattern of active genes revealed through genome-wide chromatin analysis of a higher eukaryote. Genes & development. 18:1263–1271.

Solomon, M.J., P.L. Larsen, and A. Varshavsky. 1988. Mapping protein-DNA interactions in vivo with formaldehyde: evidence that histone H4 is retained on a highly transcribed gene. Cell. 53:937–947.

Solomon, M.J., and A. Varshavsky. 1985. Formaldehyde-mediated DNA-protein crosslinking: a probe for in vivo chromatin structures. Proc Natl Acad Sci U S A. 82:6470–6474.

Spencer, V.A., J.M. Sun, L. Li, and J.R. Davie. 2003. Chromatin immunoprecipitation: a tool for studying histone acetylation and transcription factor binding. Methods. 31:67–75.

Strenkert, D., S. Schmollinger, and M. Schroda. 2011. Protocol: methodology for chromatin immunoprecipitation (ChIP) in Chlamydomonas reinhardtii. Plant Methods. 7:35.

Wang, X.F., A.A. Elling, X.Y. Li, N. Li, Z.Y. Peng, G.M. He, H. Sun, Y.J. Qi, X.S. Liu, and X.W. Deng. 2009. Genome-Wide and Organ-Specific Landscapes of Epigenetic Modifications and Their Relationships to mRNA and Small RNA Transcriptomes in Maize. The Plant cell. 21:1053–1069.

Workman, J.L., and R.E. Kingston. 1998. Alteration of nucleosome structure as a mechanism of transcriptional regulation. Annu Rev Biochem. 67:545–579.

Zhang, X., Y.V. Bernatavichute, S. Cokus, M. Pellegrini, and S.E. Jacobsen. 2009. Genome-wide analysis of mono-, di-and trimethylation of histone H3 lysine 4 in Arabidopsis thaliana. Genome biology. 10:R62.

Zhang, Y., T. Liu, C.A. Meyer, J. Eeckhoute, D.S. Johnson, B.E. Bernstein, C. Nusbaum, R.M. Myers, M. Brown, W. Li, and X.S. Liu. 2008. Model-based analysis of ChIP-Seq (MACS). Genome biology. 9:R137.

Zong, W., X.C. Zhong, J. You, and L.Z. Xiong. 2013. Genome-wide profiling of histone H3K4-tri-methylation and gene expression in rice under drought stress. Plant molecular biology. 81:175–188.

