## Supplemental Figure 1 for "An optimized ChIP-Seq framework for profiling of histone modifications in *Chromochloris zofingiensis*"

```
Cz05g04150.t1      MARTKQTARKSTGGKAPRKQLATKAARKSAPATGGVKKPHRYRPGTVALREIRKYQKSTE 60
CAB02546.1         MARTKQTARKSTGGKAPRKQLATKAARKSAPATGGVKKPHRYRPGTVALREIRRYQKSTE 60
                   *****************************************************:******

Cz05g04150.t1      LLIRKLPFQRLVREIAQDFKTDLRFQSSAVLALQEAAEAYLVGLFEDTNLCAIHAKRVTI 120
CAB02546.1         LLIRKLPFQRLVREIAQDFKTDLRFQSSAVMALQEACEATLVGLFEDTNLCAIHAKRVTI 120
                   ******************************:*****.** ********************

Cz05g04150.t1      MPKDIQLARRIRGERA*    136
CAB02546.1         MPKDIQLARRIRGERA-    136
                   ****************
```

**Supplemental Figure 1. Histone H3 is a highly conserved protein between Chromochloris and Human.** Multiple sequence alignment was performed using Clustal Omega. Multiple sequence alignment of Histone H3: *Chromochloris zofingiensis* (Cz05g04150), *Homo sapiens* (CAB02546.1). Under the alignment, asterisks indicate identical residues between both organisms.
